# Efficient implementation of penalized regression for genetic risk prediction

**DOI:** 10.1101/403337

**Authors:** Florian Privé, Hugues Aschard, Michael G.B. Blum

## Abstract

Polygenic Risk Scores (PRS) consist in combining the information across many single-nucleotide polymorphisms (SNPs) in a score reflecting the genetic risk of developing a disease. PRS might have a major impact on public health, possibly allowing for screening campaigns to identify high-genetic risk individuals for a given disease. The “Clumping+Thresholding” (C+T) approach is the most common method to derive PRS. C+T uses only univariate genome-wide association studies (GWAS) summary statistics, which makes it fast and easy to use. However, previous work showed that jointly estimating SNP effects for computing PRS has the potential to significantly improve the predictive performance of PRS as compared to C+T.

In this paper, we present an efficient method to jointly estimate SNP effects, allowing for practical application of penalized logistic regression (PLR) on modern datasets including hundreds of thousands of individuals. Moreover, our implementation of PLR directly includes automatic choices for hyper-parameters. The choice of hyper-parameters for a predictive model is very important since it can dramatically impact its predictive performance. As an example, AUC values range from less than 60% to 90% in a model with 30 causal SNPs, depending on the p-value threshold in C+T.

We compare the performance of PLR, C+T and a derivation of random forests using both real and simulated data. PLR consistently achieves higher predictive performance than the two other methods while being as fast as C+T. We find that improvement in predictive performance is more pronounced when there are few effects located in nearby genomic regions with correlated SNPs; for instance, AUC values increase from 83% with the best prediction of C+T to 92.5% with PLR. We confirm these results in a data analysis of a case-control study for celiac disease where PLR and the standard C+T method achieve AUC of 89% and of 82.5%.

In conclusion, our study demonstrates that penalized logistic regression can achieve more discriminative polygenic risk scores, while being applicable to large-scale individual-level data thanks to the implementation we provide in the R package bigstatsr.

## 1 Introduction

Polygenic Risk Scores (PRS) consist in combining the information across many single-nucleotide polymorphisms (SNPs) in a score reflecting the genetic risk of developing a disease. PRS are useful for genetic epidemiology when testing the polygenicity of one disease and finding a common genetic contribution between two diseases (Purcell *et al.* 2009). Personalized medicine is another major application of PRS. Personalized medicine envisions to use PRS in screening campaigns in order to identify high-risk individuals for a given disease (Chatterjee *et al.* 2016). As an example of practical application, targeting screening to men at higher polygenic risk could reduce the problem of overdiagnosis and lead to a better benefit-to-harm balance in screening for prostate cancer (Pashayan *et al.* 2015). Yet, PRS would have to show a high discriminative power between cases and controls in order to be used for helping in the diagnosis of diseases. For screening high-risk individuals and for presymptomatic diagnosis of the general population, it is suggested that the AUC must be greater than 75% and 99% respectively (Janssens *et al.* 2007).

Several methods have been developed to predict disease status, or more generally any phenotype, based on SNP information. A commonly used method often called “P+T” or “C+T” (which stands for “Clumping and Thresholding”) is used to derive PRS from results of Genome-Wide Association Studies (GWAS) (Chatterjee *et al.* 2013; Dudbridge 2013; Evans *et al.* 2009; Purcell *et al.* 2009; Wray *et al.* 2007). This technique uses GWAS summary statistics only, allowing for a fast implementation of C+T. However, C+T also has several limitations; for instance, previous studies have shown that predictive performance of C+T is very sensitive to the threshold of inclusion of SNPs, depending on the disease architecture (Ware *et al.* 2017). Linear Mixed-Models (LMMs) are another widely-used method in fields such as plant and animal breeding or for predicting highly heritable quantitative human phenotypes such as height (Lello *et al.* 2017; Yang *et al.* 2010). Yet, models resulting from LMM, known e.g. as “gBLUP”, are not optimal for predicting disease status based on genotypes (Abraham *et al.* 2013). Moreover, these methods and their derivatives are often computationally very demanding, both in terms of memory and time required, which makes them unlikely to be used for prediction on very large datasets (Golan and Rosset 2014; Maier *et al.* 2015; Speed and Balding 2014; Zhou *et al.* 2013). Finally, statistical learning methods have also been used to derive PRS for complex human diseases by jointly estimating SNP effects. Such methods include joint logistic regression, Support Vector Machine (SVM) and random forests (Abraham *et al.* 2012, 2014; *Botta et al.* 2014; Okser *et al.* 2014; Wei *et al.* 2009).

We recently developed two R packages, bigstatsr and bigsnpr, for efficiently analyzing large-scale genome-wide data (Privé *et al.* 2018). Package bigstatsr now includes an efficient algorithm with a new implementation for computing sparse linear and logistic regressions on huge datasets as large as the UK Biobank (Bycroft *et al.* 2017). In this paper, we present a comprehensive comparative study of our implementation of penalized logistic regression (PLR) against the C+T method and the T-Trees algorithm, a derivation of random forests that has shown high predictive performance (Botta *et al.* 2014). In this comparison, we do not include any LMM method for the reasons mentioned before and do not include any SVM method because it is expected to give similar results to logistic regression (Abraham *et al.* 2012). For C+T, we report results for a large grid of hyper-parameters. For PLR, the choice of hyper-parameters is included in the algorithm so that we report only one model for each simulation. We also use a modified version of PLR in order to capture not only linear effects, but also recessive and dominant effects.

To perform simulations, we use real genotype data and simulate new phenotypes. In order to make our comparison as comprehensive as possible, we compare different disease architectures by varying the number, size and location of causal effects as well as the disease heritability. We also compare two different models for simulating phenotypes, one with additive effects only, and one that combines additive, dominant and interaction-type effects. Overall, we find that PLR consistently achieves higher predictive performance than the C+T and T-Trees methods while being as fast as C+T. This demonstrates the feasibility and relevance of this approach for PRS computation on large modern datasets.

## 2 Material and Methods

### 2.1 Genotype data

We use real genotypes of European individuals from a case-control study for celiac disease (Dubois *et al.* 2010). The composition of this dataset is presented in table S1. Details of quality control and imputation for this dataset are available in Privé *et al.* (2018). For simulations presented later, we first restrict this dataset to controls from UK in order to remove the genetic structure induced by the celiac disease status and population structure. This filtering process results in a sample of 7100 individuals (see supplementary notebook “preprocessing”). We also use this dataset for real data application, in this case keeping all 15,155 individuals (4496 cases and 10,659 controls). Both datasets contain 281,122 SNPs.

### 2.2 Simulations of phenotypes

We simulate binary phenotypes using a Liability Threshold Model (LTM) with a prevalence of 30% (Falconer 1965). We vary simulation parameters in order to match a range of genetic architectures from low to high polygenicity. This is achieved by varying the number of causal variants and their location (30, 300, or 3000 anywhere in all 22 autosomal chromosomes or 30 in the HLA region of chromosome 6), and the disease heritability *h*^2^ (50% or 80%). Liability scores are computed either from a model with additive effects only (“ADD”) or a more complex model that combines additive, dominant and interaction-type effects (“COMP”). For model “ADD”, we compute the liability score of the i-th individual

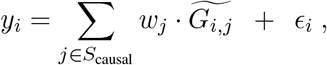

where *S*_causal_ is the set of causal SNPs, *w_j_* are weights generated from a Gaussian distribution *N*(0, *h*^2^/|*S*_causal_|) or a Laplace distribution 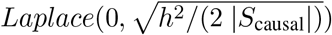, *G_i,j_* is the allele count of individual *i* for SNP *j*, 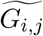 corresponds to its standardized version (zero mean and unit variance for all SNPs), and *ε* follows a Gaussian distribution *N* (0, 1 − *h*^2^). For model “COMP”, we simulate liability scores using additive, dominant and interaction-type effects (see Supplementary Materials).

We implement 3 different simulation scenarios, summarized in table 2. Scenario 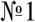 uses the whole dataset (all 22 autosomal chromosomes – 281,122 SNPs) and a training set of size 6000. It compares all methods described in section 2.4. For each combination of the remaining parameters, results are based on 100 simulations excepted when comparing PLR with T-Trees, which relies on 5 simulations only because of a much higher computational burden of T-Trees as compared to other methods. Scenario 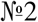 consists of 100 simulations per combination of parameters on a dataset composed of chromosome 6 only (18,941 SNPs). Reducing the number of SNPs increases the polygenicity (i.e. the proportion of causal SNPs) of the simulated models. Reducing the number of SNPs (*p*) is also equivalent to increasing the sample size (*n*) as predictive power is dependent on *n/p* (Dudbridge 2013; Vilhjálmsson *et al.* 2015). For this scenario, we use the additive model only, but continue to vary all other simulation parameters. Finally, scenario 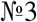 uses the whole dataset as in scenario 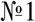 while varying the size of the training set in order to assess how the sample size affects predictive performance of methods. A total of 100 simulations per combination of parameters are run using 300 causal SNPs randomly chosen on the genome.

### 2.3 Predictive performance measures

In this study, we use two different measures of predictive accuracy. First, we use the Area Under the Receiver Operating Characteristic (ROC) Curve (AUC) (Fawcett 2006; Lusted 1971). In the case of our study, the AUC is the probability that the PRS of a case is greater than the PRS of a control. This measure indicates the extent to which we can distinguish between cases and controls using PRS. As a second measure, we also report the partial AUC for specificities between 90% and 100% (Dodd and Pepe 2003; McClish 1989). This measure is similar to the AUC, but focuses on high specificities, which is the most useful part of the ROC curve in clinical settings. When reporting AUC results of simulations, we also report maximum achievable AUC values of 84% and 94% for heritabilities of 50% and 80% respectively. These estimates are based on three different yet consistent estimations (see Supplementary Materials).

### 2.4 Methods compared

In this paper, we compare three different types of methods: the C+T method, T-Trees and penalized logistic regression (PLR).

The C+T (Clumping + Thresholding) method directly derives a Polygenic Risk Score (PRS) from the results of Genome-Wide Associations Studies (GWAS). In GWAS, a coefficient of regression (i.e. the estimated effect size 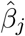) is learned independently for each SNP *j* along with a corresponding p-value *p_j_*. The SNPs are first clumped (C) so that there remain only loci that are weakly correlated with one another (this set of SNPs is denoted *S*_clumping_). Then, thresholding (T) consists in removing SNPs with p-values larger than a user-defined threshold *p_T_*. Finally, the PRS for individual *i* is defined as the sum of allele counts of the remaining SNPs weighted by the corresponding effect coefficients

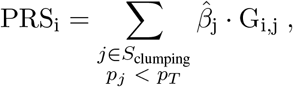

where 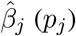 are the effect sizes (p-values) learned from the GWAS. In this study, we mostly report scores for a clumping threshold at *r*^2^ > 0.2 within regions of 500kb, but we also investigate thresholds of 0.05 and 0.8. We report three different scores of prediction: one including all the SNPs remaining after clumping (denoted “C+T-all”), one including only the SNPs remaining after clumping and that have a p-value under the GWAS threshold of significance (*p* < 5 · 10^−8^, “C+T-stringent”), and one that maximizes the AUC (“C+T-max”) for 102 p-value thresholds between 1 and 10^−100^ (Table S2). As we report the optimal threshold based on the test set, the AUC for “C+T-max” is an upper bound of the AUC for the C+T method.

T-Trees (*Trees inside Trees*) is an algorithm derived from random forests (Breiman 2001) that takes into account the correlation structure among the genetic markers implied by linkage disequilibrium in GWAS data (Botta *et al.* 2014). We use the same parameters as reported in Table 4 of Botta *et al.* (2014), except that we use 100 trees instead of 1000. Using 1000 trees provides a minimal increase of AUC while requiring a disproportionately long processing time (e.g. AUC of 81.5% instead of 81%, data not shown).

Finally, for penalized logistic regression (PLR), we find regression coefficients *β*_0_ and *β* that minimize the following regularized loss function

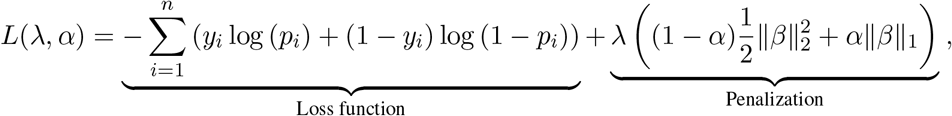

where 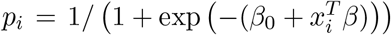, *x* is denoting the genotypes and covariables (e.g. principal components), *y* is the disease status to predict, *λ* and *α* are two regularization hyper-parameters that need to be chosen. Different regularizations can be used to prevent overfitting, among other benefits: the L2-regularization (“ridge”, Hoerl and Kennard (1970)) shrinks coefficients and is ideal if there are many predictors drawn from a Gaussian distribution (corresponds to *α* = 0 in the previous equation); the L1-regularization (“lasso”, Tibshirani (1996)) forces some of the coefficients to be equal to zero and can be used as a means of variable selection, leading to sparse models (corresponds to *α* = 1); the L1- and L2-regularization (“elastic-net”, Zou and Hastie (2005)) is a compromise between the two previous penalties and is particularly useful in the *p ≫ n* situation (*p* is the number of SNPs), or any situation involving many correlated predictors (corresponds to 0 < *α* < 1) (Friedman *et al.* 2010). In this study, we use an embedded grid search over *α* ∈ {1, 0.5, 0.05, 0.001}.

To fit this penalized logistic regression, we use an efficient algorithm (Friedman *et al.* 2010; Tibshirani *et al.* 2012; Zeng *et al.* 2017) from which we derived our own implementation in R package bigstatsr. This type of algorithm builds predictions for many values of *λ*, which is called a “regularization path”. To obtain an algorithm free of the choice of this hyper-parameter *λ*, we developed a procedure that we call Cross-Model Selection and Averaging (CMSA, figure S1). Because of L1-regularization, the resulting vectors of coefficients are sparse and can be used to make a PRS based on a *linear* combination of allele counts. We refer to this method as “PLR” in the results section.

To capture recessive and dominant effects on top of additive effects in PLR, we use simple feature engineering: we construct a separate dataset with 3 times as many variables as the initial one. For each SNP variable, we add two more variables coding for recessive and dominant effects: one variable is coded 1 if homozygous variant and 0 otherwise, and the other is coded 0 for homozygous referent and 1 otherwise. We then apply our PLR implementation to this dataset with 3 times as many variables as the initial one; we refer to this method as “PLR3” in the rest of the paper.

### 2.5 Evaluating predictive performance for Celiac data

We use Monte Carlo cross-validation to compute AUC, partial AUC, the number of predictors and execution time for the original Celiac dataset with the observed case-control status: we randomly split 100 times the dataset in a training set of 12,000 indiduals and a test set composed of the remaining 3155 individuals.

## 3 Results

### 3.1 Joint estimation improves predictive performance

We compared penalized logistic regression (PLR) with the C+T method using simulations of scenario 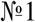 (Table 2). When simulating a model with 30 causal SNPs and an heritability of 80%, PLR provides AUC of 93%, nearly reaching the maximum achievable AUC of 94% for this setting (Figure 1). Moreover, PLR consistently provides higher predictive performance than C+T across all scenarios we considered, excepted in some cases of high polygenicity or small sample size where all methods perform poorly (AUC values below 60% − figures 3 and S3). PLR provides particularly higher predictive performance than C+T when there are correlations between predictors, i.e. when we choose causal SNPs to be in the HLA region. In this situation, the mean AUC reaches 92.5% for PLR and 84% for “C+T-max” (Figure 1). Note that, for the simulations, we do not report results in terms of partial AUC because partial AUC values have a Spearman correlation of 98% with the AUC results for all methods (Figure S2).

**Figure 1:**
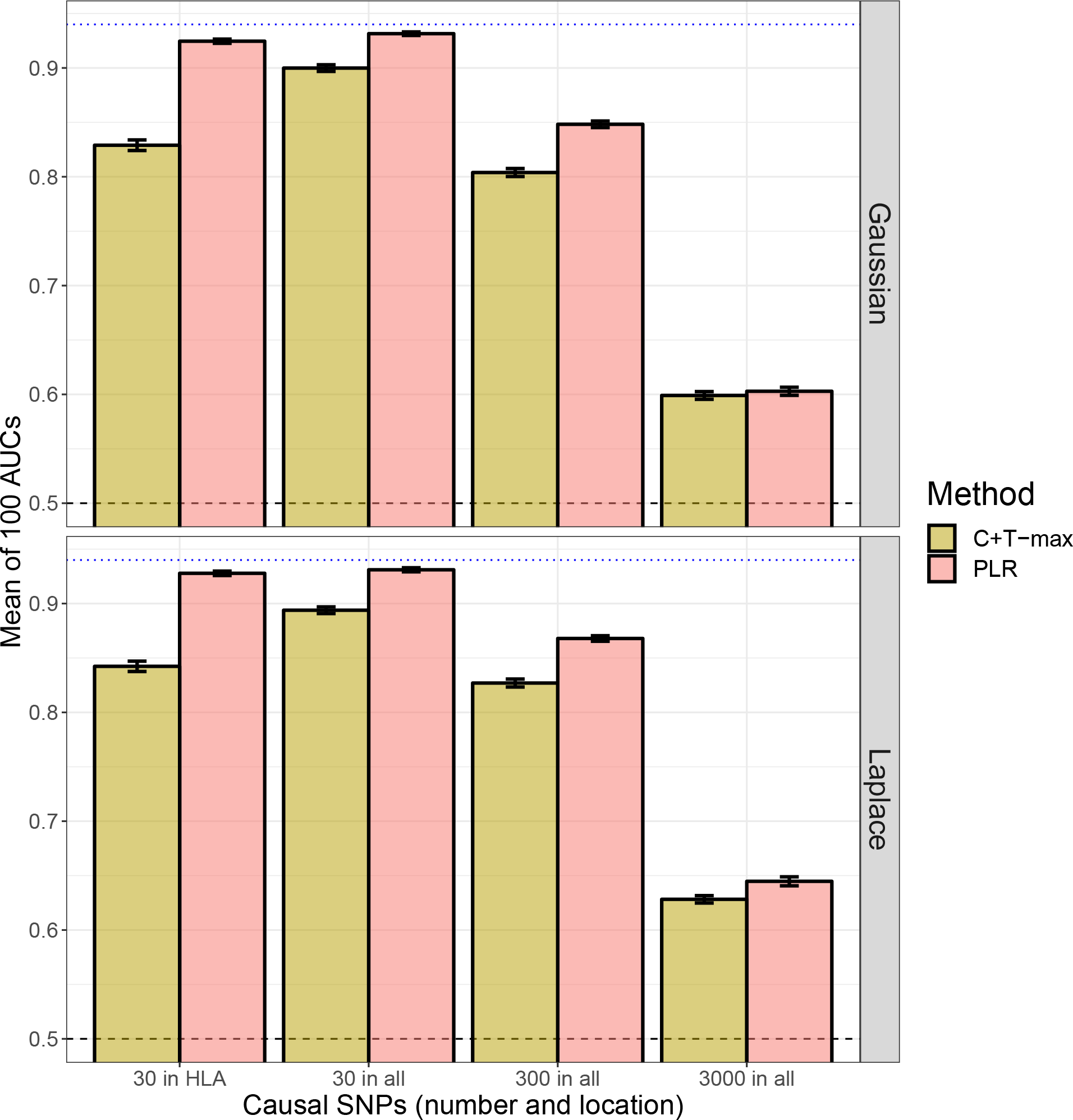
Main comparison of C+T and PLR in scenario 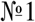 for model “ADD” and an heritability of 80%. Mean AUC over 100 simulations for PLR and the maximum AUC reported with “C+T-max”. Upper (lower) panel is presenting results for effets following a Gaussian (Laplace) distribution. Error bars are representing ±2SD of 10^5^ non-parametric bootstrap of the mean AUC. The blue dotted line represents the maximum achievable AUC.

**Figure 2:**
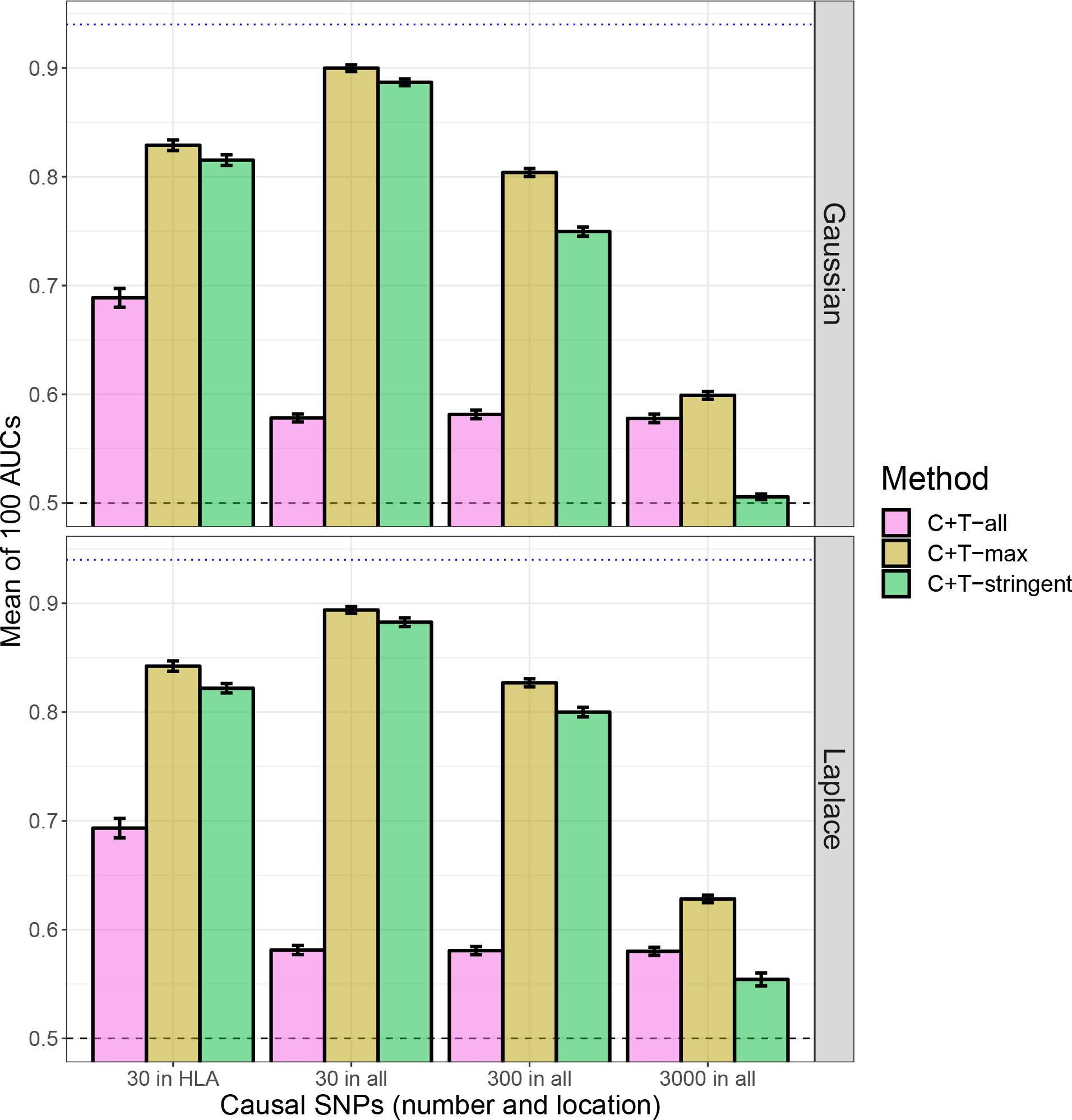
Comparison of three different p-value thresholds used in the C+T method in scenario 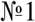 for model “ADD” and an heritability of 80%. Mean AUC over 100 simulations. Upper (lower) panel is presenting results for effets following a Gaussian (Laplace) distribution. Error bars are representing ±2SD of 10^5^ non-parametric bootstrap of the mean AUC. The blue dotted line represents the maximum achievable AUC.

**Figure 3:**
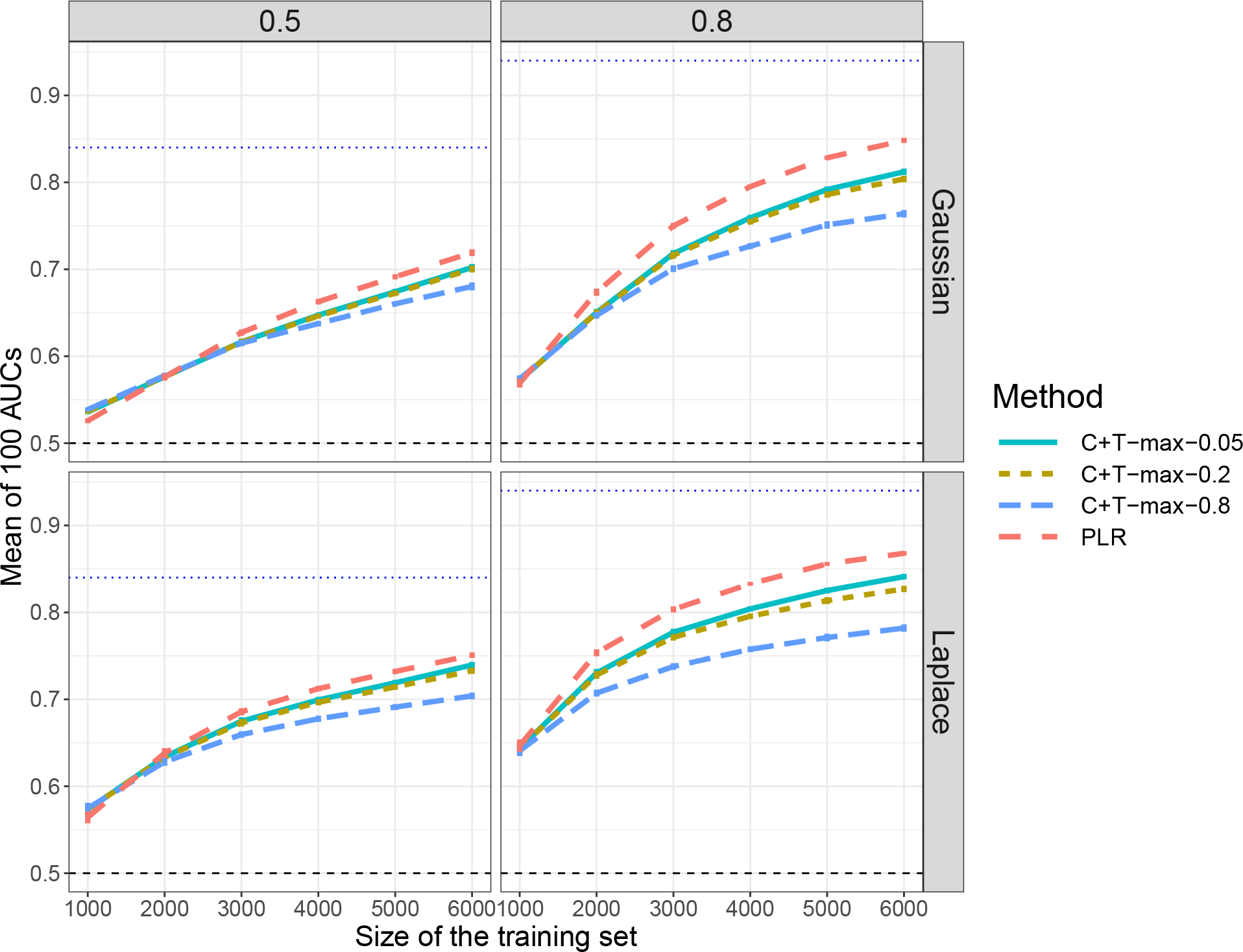
Comparison of methods when varying sample size in scenario 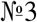 for model “ADD” with 300 causal SNPs sampled anywhere on the genome. Mean AUC over 100 simulations for the maximum values of C+T for three different *r*^2^ thresholds (0.05, 0.2 and 0.8) and PLR as a function of the training size. Upper (lower) panels are presenting results for effets following a Gaussian (Laplace) distribution and left (right) panels are presenting results for an heritability of 0.5 (0.8). Error bars are representing ±2SD of 10^5^ non-parametric bootstrap of the mean AUC. The blue dotted line represents the maximum achievable AUC.

### 3.2 Importance of hyper-parameters

In practice, a particular value of the threshold of inclusion of SNPs should be chosen for the C+T method and this choice can dramatically impact the predictive performance of C+T. For example, in a model with 30 causal SNPs, AUC ranges from less than 60% when using all SNPs passing clumping to 90% *if* choosing the optimal p-value threshold (Figures 2 and S4).

Concerning the *r*^2^ threshold of the clumping step in C+T, we mostly used the common value of 0.2. Yet, using a more stringent value of 0.05 provides higher predictive performance than using 0.2 in most of the cases we considered (Figures S5, 3 and S6)

Our implementation of PLR that automatically chooses hyper-parameter *λ* provides similar predictive performance than the best predictive performance of 100 models corresponding to different values of *λ* (Figure S10).

### 3.3 Non-linear effects

We tested the T-Trees method in scenario 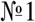. As compared to PLR, T-Trees perform worse in terms of predictive ability, while taking much longer to run (Figure S7). Even for simulations with model “COMP” in which there are dominant and interaction-type effects that T-Trees should be able to handle, AUC is still lower when using T-Trees than when using PLR (Figure S7).

We also compared the two penalized logistic regressions in scenario 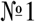: PLR versus PLR3 that uses additional features (variables) coding for recessive and dominant effects. Predictive performance of PLR3 are nearly as good as PLR when there are additive effects only (differences of AUC are always smaller than 2%) and can lead to significantly greater results when there are also dominant and interactions effects (Figures S8 and S9). For model “COMP”, PLR3 provides AUC values at least 3.5% higher than PLR, excepted when there are 3000 causal SNPs. Yet, PLR3 takes 2-3 times as much time to run and requires 3 times as much disk storage as PLR.

### 3.4 Simulations varying number of SNPs and training size

First, when reproducing simulations of scenario 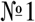 using chromosome 6 only (scenario 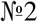), the predictive performance of PLR always increase (Figure S6). There is a particularly large increase when simulating 3000 causal SNPs: AUC from PLR increases from 60% to nearly 80% for Gaussian effects and a disease heritability of 80%. On the contrary, when simulating only 30 or 300 causal SNPs with the corresponding dataset, AUC of “C+T-max” does not increase, and even decreases for an heritability of 80% (Figure S6). Secondly, when varying the training size (scenario 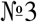), we report an increase of AUC with a larger training size, with a faster increase of AUC for PLR as compared to “C+T-max” (Figure 3).

### 3.5 Polygenic scores for the celiac disease

Joint logistic regressions also provide higher AUC values for the Celiac data: 88.7% with PLR and 89.1% with PLR3 as compared to 82.5% with “C+T-max”. The relative increase in partial AUC, for specificities larger than 90%, is even larger (42% and 47%) with partial AUC values of 0.0411, 0.0426 and 0.0289 obtained with PLR, PLR3 and “C+T-max”, respectively. Moreover, logistic regressions use less predictors, respectively 1570, 2260 and 8360 (Table 1, figure 4 and supplementary notebook “results-celiac”). In terms of computation time, we show that PLR, while learning jointly on all SNPs at once and testing four different values for hyper-parameter *α*, is almost as fast as the C+T method (190 vs 130 seconds), and PLR3 takes less than twice as long as PLR (296 vs 190 seconds).

**Table 1:** Results for the real Celiac dataset. The results are averaged over 100 runs where the training step is randomly composed of 12,000 individuals. In the parentheses is reported the standard deviation of 10^5^ bootstrap samples of the mean of the corresponding variable. Results are reported with 3 significant digits.

| Method | AUC | pAUC | # predictors | Execution time (s) |
| --- | --- | --- | --- | --- |
| C+T-max | 0.825 (0.000664) | 0.0289 (0.000187) | 8360 (744) | 130 (0.143) |
| PLR | 0.887 (0.00061) | 0.0411 (0.000224) | 1570 (46.4) | 190 (1.21) |
| PLR3 | 0.891 (0.000628) | 0.0426 (0.000219) | 2260 (56.1) | 296 (2.03) |

**Figure 4:**
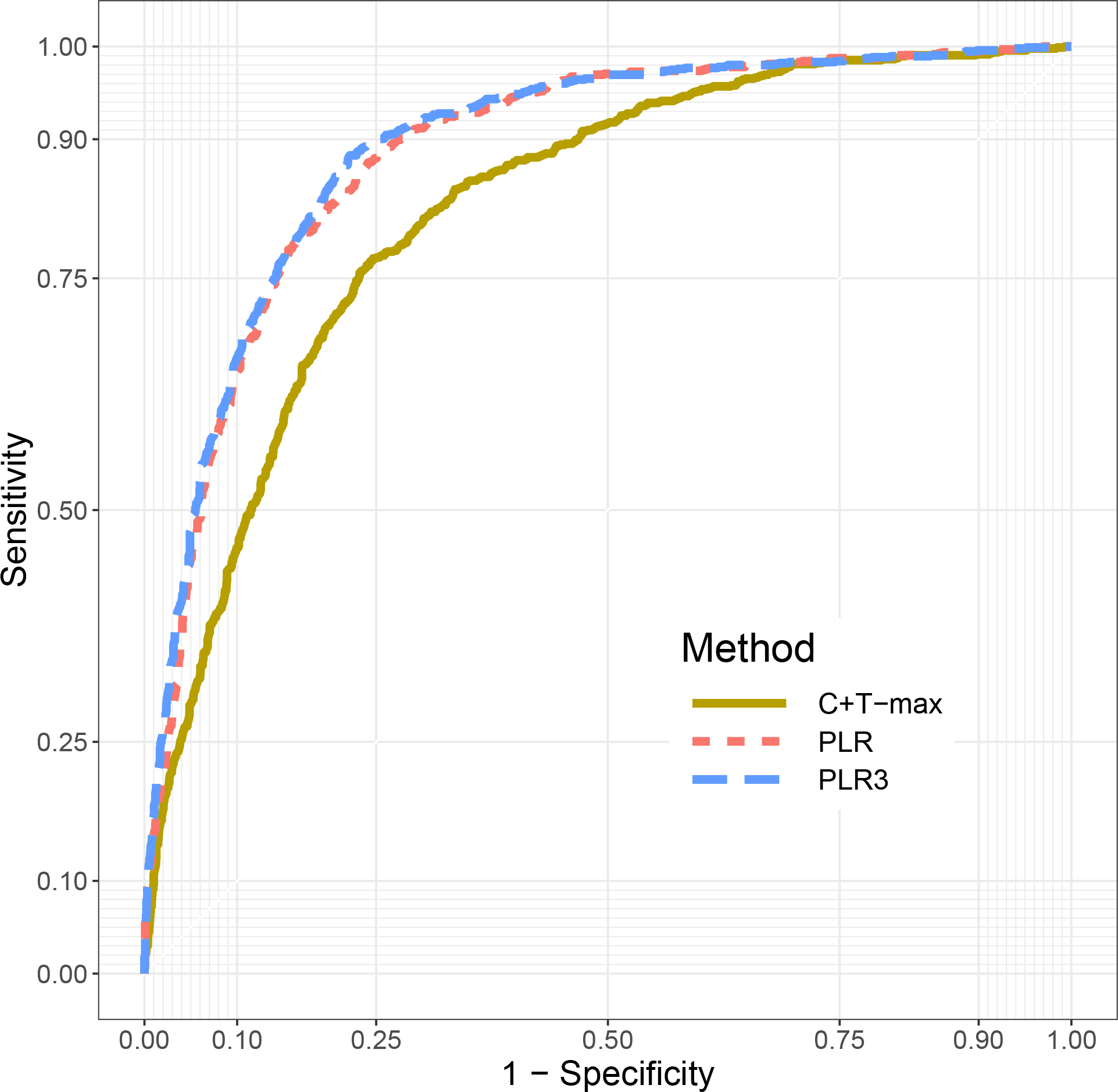
ROC Curves for C+T, PLR and PLR3 for the celiac disease dataset. Models were trained using 12,000 individuals. These are results projecting these models on the remaining 3155 individuals. The figure is plotted using R package plotROC (Sachs *et al.* 2017).

## 4 Discussion

### 4.1 Joint estimation improves predictive performance

In this comparative study, we present a computationally efficient implementation of penalized logistic regression (PLR). This model can be used to build polygenic risk scores based on very large individual-level SNP datasets such as the UK biobank (Bycroft *et al.* 2017). In agreement with previous work (Abraham *et al.* 2013), we show that jointly estimating SNP effects has the potential to substantially improve predictive performance as compared to the standard C+T approach in which SNP effects are learned independently. PLR always outperform the C+T method, excepted in some highly underpowered cases, and the benefits of using PLR are more pronounced with an increasing sample size or when causal SNPs are correlated with one another.

### 4.2 Importance of hyper-parameters

The choice of hyper-parameter values is very important since it can greatly impact method performance. In the C+T method, there are two main hyper-parameters: the *r*^2^ and the *p_T_* thresholds that control how stringent are the clumping and thresholding steps, respectively. The choice of the *r*^2^ threshold of the clumping step is important. Indeed, on the one hand, choosing a low value for this threshold may discard informative SNPs that are correlated. Yet, on the other hand, when choosing a high value for this threshold, too much redundant information would be included in the model, which would add some noise to the PRS. Based on the simulations, we find that using a stringent threshold (*r*^2^ = 0.05) leads to higher predictive performance, even when causal SNPs are correlated. It means that, in most cases, avoiding redundant information is more important than including all causal SNPs. The choice of the *p_T_* threshold is also very important as it can greatly impact the predictive performance of the C+T method, which we confirm in this study (Ware *et al.* 2017). In this paper, we reported the maximum AUC of 102 different p-value thresholds, a threshold that should normally be learned on the training set only. To our knowledge, there is no clear standard on how to choose these two critical hyper-parameters for C+T.

On the contrary, for the penalized logistic regression presented here, we developed an automatic procedure called Cross-Model Selection and Averaging (CMSA) that releases investigators from the burden of choosing hyper-parameter *λ* that accounts for the amount of regularization used in the model. Not only this procedure provides near-optimal results, but it also accelerates the model training thanks to the development of an early stopping criterion. Usually, cross-validation is used to choose hyper-parameter values and then the model is trained again with these particular hyper-parameter values (Hastie *et al.* 2008; Wei *et al.* 2013). Yet, performing cross-validation and retraining the model is computationally demanding; CMSA offers a less burdensome alternative. Concerning hyper-parameter *α* that accounts for the relative importance of the L1 and L2 regularizations, we use a grid search directly embedded in the CMSA procedure.

### 4.3 Non-linear effects

We also explored how to capture non-linear effects. For this, we introduced a simple feature engineering technique that enables PLR to detect and learn not only additive effects, but also dominant and recessive effects. This technique improves the predictive performance of PLR when there are some non-linear effects in the simulations, while providing nearly the same predictive performance when there are additive effects only. Moreover, it also improves predictive performance for the celiac disease.

Yet, this approach is not able to detect interaction-type effects. In order to capture interaction-type effects, we tested T-Trees, a method that is able to exploit SNP correlations and interactions thanks to special decision trees (Botta *et al.* 2014). However, predictive performance of T-Trees are consistently lower than with penalized logistic regression, even when simulating a model with dominant and interaction-type effects that T-Trees should be able to handle.

### 4.4 Limitations

Our approach has one major limitation: the main advantage of the C+T method is its direct applicability to summary statistics, allowing to leverage the largest GWAS results to date, even when individual cohort data cannot be merged because of practical or ethical reasons (e.g. consortium data including many cohorts). As of today, the proposed penalized logistic regression does not allow for the analysis of summary data, but this represents an important future direction of our work. The current version is of particular interest for the analysis of modern individual-level datasets including hundreds of thousands of individuals.

Finally, in this comparative study, we did not consider the problem of population structure (Márquez-Luna *et al.* 2017; Martin *et al.* 2017; Vilhjálmsson *et al.* 2015) and also did not consider non-genetic data such as environmental and clinical data (Dey *et al.* 2013; Van Vliet *et al.* 2012).

### 4.5 Conclusion

In this comparative study, we have presented a computationally efficient implementation of penalized logistic regression that can be used to predict disease status based on genotypes. Note that a similar penalized linear regression is also available in our software. Our approach solves the dramatic computational burden faced by standard implementations, thus allowing for the analysis of large-scale datasets such as the UK biobank (Bycroft *et al.* 2017).

We also demonstrated in simulations that our implementation of penalized regressions remains highly effective over a broad range of disease architectures. It can be appropriate for predicting autoimmune diseases with a few strong effects (e.g. celiac disease) as well as highly polygenic traits (e.g. standing height). Finally, note that these models could also be used to predict phenotypes based on other omics data since the implementation is not specific to genotype data.

**Table 2:** Summary of all simulations. Where there is symbol ‘−’ in a box, it means that the parameters are the same as the ones in the upper box.

| Numero of scenario | Dataset | Size of training set | Causal SNPs (number and location) | Distribution of effects | Heritability | Simulation model | Methods |
| --- | --- | --- | --- | --- | --- | --- | --- |
| 1 | All 22 chromosomes | 6000 | 30 in HLA<br>30 in all<br>300 in all<br>3000 in all | Gaussian<br>Laplace | 0.5<br>0.8 | ADD<br>COMP | C+T<br>PLR<br>PLR3<br>(T-Trees) |
| 2 | Chromosome 6 only | - | - | - | - | ADD | C+T<br>PLR |
| 3 | All 22 chromosomes | 1000<br>2000<br>3000<br>4000<br>5000 | 300 in all | - | - | - | - |

## Description of Supplemental Data

Supplemental Data include a PDF with two sections of methods, two tables and ten figures. Supplemental Data also include six HTML R notebooks including all code and results used in this paper, for reproducibility purposes, and available at https://figshare.com/articles/code/7178750.

## Declaration of Interests

The authors declare no competing interests.

## Acknowledgements

Authors acknowledge LabEx PERSYVAL-Lab (ANR-11-LABX-0025-01). Authors also acknowledge the Grenoble Alpes Data Institute that is supported by the French National Research Agency under the “Investissements d’avenir” program (ANR-15-IDEX-02). We are also grateful to Félix Balazard for useful discussions about T-Trees, and to Yaohui Zeng for useful discussions about R package biglasso.

## Web Resources

Results of simulations are available at https://figshare.com/articles/results_zip/7126964. A tutorial on how to start with R packages bigstatsr and bigsnpr is available at https://privefl.github.io/bigsnpr/articles/demo.html. The two R packages are available on GitHub.

